# The targeted cytosolic degradation of class I histone deacetylases is essential for efficient alphaherpesvirus replication

**DOI:** 10.64898/2026.01.02.697337

**Authors:** Sheng-Li Ming, Meng-Hua Du, Jia-Ming Yang, Ya-Di Guo, Jia-Jia Pan, Wei-Fei Lu, Jiang Wang, Lei Zeng, Bei-Bei Chu

**Author notes:** These authors contributed equally (Sheng-Li Ming, Meng-Hua Du, Jia-Ming Yang). Correspondence (L. Zeng); (B.-B. Chu).

## Abstract

Viral infection triggers a robust DNA damage response (DDR), reshaping the host chromatin landscape to facilitate viral replication. Here, we uncover a novel mechanism by which alphaherpesviruses exploit the DDR pathway. We demonstrated that herpes simplex virus 1 (HSV-1) and pseudorabies virus (PRV) induced selective degradation of class I histone deacetylases (HDAC1/2), leading to histone hyperacetylation and subsequent DDR activation. Strikingly, viral infection promoted nuclear export of HDAC1/2, followed by MDM2-mediated K63-linked polyubiquitination and proteasomal degradation in the cytoplasm. Pharmacological inhibition of either DDR signaling or HDAC1/2 nuclear export significantly affected viral replication *in vitro* and *in vivo*. Our findings reveal a unique viral strategy to hijack host epigenetic regulation for efficient replication and identify potential therapeutic targets for alphaherpesvirus infections.

## Introduction

HSV-1 is a widespread neurotropic virus that infects more than two-thirds of the global population under the age of 50 [1]. After initial infection of mucosal epithelia and neuronal tissues, HSV-1 establishes lifelong latency within sensory ganglia [2]. Clinically, HSV-1 infection can present as oral or genital lesions and, in severe instances, may progress to herpes simplex encephalitis–a potentially fatal inflammation of the brain [3, 4]. Emerging evidence suggests a plausible association between HSV-1 and neurodegenerative conditions such as Alzheimer’s disease, linking chronic neuroinflammation and neuronal impairment to persistent viral infection [5, 6]. To sustain lifelong infection, HSV-1 employs sophisticated strategies to evade host immune defenses and maintain latent reservoirs.

Epigenetic regulation plays a critical role in governing HSV-1 latency and reactivation [7, 8]. During latency, the virus coopts host histone deacetylases (HDACs) to enforce viral genomic silencing through chromatin condensation [9]. Although HDAC1 and HDAC2 are central to chromatin stability and DNA damage response (DDR), their roles during alphaherpesvirus infection appear complex and context-dependent [10]. While these enzymes contribute to viral repression during latency [11], their potential involvement in promoting lytic replication–whether through degradation or functional inactivation–has remained inadequately explored.

Histone acetylation, a pivotal epigenetic modification that modulates chromatin accessibility and transcriptional activity, is dynamically regulated by histone acetyltransferases (HATs), which add acetyl groups to activate gene expression, and HDACs, which remove these groups to enforce repression [12, 13]. The HDAC family is categorized into four classes: Class I (HDAC1, 2, 3, 8), Class IIa/b, Class III (sirtuins), and Class IV (HDAC11) [14]. Among these, Class I HDACs–particularly HDAC1 and HDAC2–are indispensable for chromatin remodeling, genomic integrity, and immune modulation [15-18]. These enzymes contribute to host antiviral defense by fine-tuning the expression of immune-related genes. Nevertheless, the mechanisms through which HSV-1 overcomes this transcriptional repression remain incompletely understood [18].

HDAC1 serves as a key regulator of chromatin dynamics and innate immune signaling, interfacing with critical pathways such as NF-κB, JAK-STAT, and Toll-like receptor cascades to shape antiviral responses [19-22]. For example, in lung epithelial cells, HDAC1 facilitates STAT1 phosphorylation and enhances interferon-stimulated gene activation, thereby restricting influenza A virus replication [23]. This underscores the dual function of HDAC1 in both epigenetic control and immune regulation. However, the specific strategies used by HSV-1 to manipulate HDAC1/2–particularly through ubiquitin-mediated degradation–to circumvent host immunity and enhance viral replication have not been clearly defined.

This study reveals that HSV-1 induces the degradation of HDAC1/2 via an MDM2-dependent ubiquitination mechanism, resulting in elevated histone acetylation, chromatin relaxation, and activation of DNA damage response pathways that collectively enhance viral replication. These findings offer novel insights into the epigenetic subversion strategies employed by HSV-1 and underscore the therapeutic potential of targeting the MDM2–HDAC axis in the treatment of herpesvirus infections.

## Materials and Methods

### Mice

Female C57BL/6J mice (6–8 weeks old) were purchased from the Experimental Animal Center of Zhengzhou University (Zhengzhou, China) and housed in specific-pathogen-free facilities under controlled environmental conditions, including a 12-hour light-dark cycle and a temperature of 22°C. For *in vivo* infection studies, mice were anesthetized with isoflurane prior to intranasal inoculation with HSV-1, at a dose of 1 × 10^6^ PFU per mouse in a total volume of 30 μL PBS. Clinical signs–including ruffled fur, hunched posture, reduced mobility, and body weight loss 15%–were monitored daily. On day 15 post-infection, mice were humanely euthanized by intravenous injection of sodium pentobarbital (90 mg/kg), following the “Guidelines for the Euthanasia of Laboratory Animals”. Immediately thereafter, tissues–including liver, spleen, lung, brain, and draining cervical lymph nodes–and serum were harvested under sterile conditions. All tissues were snap-frozen in liquid nitrogen within 60 seconds of excision and stored at −80°C until further analysis. Samples were processed for Western blotting. All procedures were in accordance with ethical guidelines.

### Reagents and plasmids

Reagents were sourced as follows: Leptomycin B (HY-16909), MG-132 (HY-13259), 3-MA (HY-19312), cycloheximide (HY-12320), and chloroquine (HY-17589A) were purchased from MedChemExpress; berzosertib (S7102) was purchased from Selleck; DMSO (W387520) was purchased from Sigma-Aldrich. Unless otherwise specified, all inhibitors were added at 1-hour post-infection (hpi), after completion of viral adsorption and entry, to avoid interference with early viral processes. Antibodies against CHK1 (25887-1-AP), CHK2 (13954-1-AP), RAD51 (14961-1-AP), β-actin (66009-1-lg), P53 (10442-1-AP), HDAC1 (10197-1-AP), HDAC2 (12922-3-AP), HDAC4 (17449-1-AP), HDAC6 (12834-1-AP), HDAC11 (67949-1-Ig), and EGFP (50430-2-AP) were purchased from Proteintech; antibodies against p-P53 (9286), p-ATM (13050), ATM (2873), ATR (13934), p-ATR (2853), p-CHK1 (12302), p-CHK2 (2197), γ-H2AX (80312), H3 (4499), H4K8ac (2594), H4K12ac (13944), H4 (13919), H2AX (7631), H3K9ac (4658), H3K27ac (8173), UB (20326), and MDM2 (86934) were purchased from Cell Signaling Technology; H3K56ac antibody (07-677) was purchased from Millipore; ICP4 antibody (ab6514) was purchased from Abcam; ICP0 antibody (sc-53070) was purchased from Santa Cruz Biotechnology; FLAG antibody (F7425) was purchased from Sigma-Aldrich; HA antibody (A00169) was purchased from GenScript. HDAC1 and HDAC2 coding sequences were amplified from HEK293 cell cDNA and cloned into p3×FLAG-CMV-10 to generate FLAG-tagged expression constructs. MDM2 coding sequence was amplified and inserted into pEGFP-C1 to produce EGFP-MDM2. Site-directed mutagenesis was performed using the QuikChange Site-Directed Mutagenesis Kit (Agilent Technologies, 200523) per the manufacturer’s instructions to generate: FLAG-HDAC1 K8R, FLAG-HDAC1 K10R, FLAG-HDAC1 K31R, FLAG-HDAC1 K50R, FLAG-HDAC1 K58R, FLAG-HDAC1 K66R, FLAG-HDAC1 K74R, FLAG-HDAC1 K89R, FLAG-HDAC2 K75R, FLAG-HDAC1 (aa 1–329), FLAG-HDAC1 (aa 329–482), FLAG-HDAC1 (aa 1–100), FLAG-HDAC1 (aa 100–329), and EGFP-MDM2ΔRING (aa 1–435). FLAG-HDAC1 ΔNES (residues 207–216 deleted), in which the deletion encompasses the CRM1-dependent nuclear export signal as predicted by NESmapper. HA-tagged ubiquitin constructs [HA-UB (WT), HA-UB (K48), HA-UB (K63)] were provided by Dr. Bo Zhong (College of Life Sciences, Wuhan University, China).

### Cells

Cell lines (HeLa/ATCC CL-82, Vero/ATCC CL-81, 3D4/21/ATCC CRL-2843, HEK293T/ATCC CRL-11268) were cultured in DMEM 10566–016 or RPMI 1640 (Gibco, 11965092) supplemented with 10% fetal bovine serum (FBS; Gibco, 10099141C), 1% penicillin/streptomycin and 1% L-glutamine (B540732, Sangon) at 37°C under 5% CO_2_ in a humidified incubator. For transient transfections, DNA constructs were delivered to HeLa and HEK293T cells using Lipofectamine 3000 (Invitrogen, L3000001) per the manufacturer’s protocol.

To generate shRNA-mediated knockdown cell lines, HEK293T cells were co-transfected with control or gene-specific shRNA (Table S1), packaging plasmid psPAX2, and envelope plasmid pMD2.G. Post-transfection (6 h), medium was replaced, and lentiviral particles were harvested at 48 hours. Parental cells were infected with viral supernatant and selected with puromycin to establish stable knockdown lines.

To generate siRNA-mediated knockdown cell lines, HeLa cells were transfected with either a non-targeting control siRNA or gene-specific siRNAs (Table S1) using Lipofectamine™ RNAiMAX (Invitrogen, 13778030). Transfection was performed at a confluency of approximately 30%, and target gene and protein expression levels were assessed 48 hours post-transfection.

### Viruses

HSV-1 strain F (provided by Dr. Chun-Fu Zheng, University of Calgary, Canada) and PRV-QXX were propagated per established protocols [24]. Viral titers were determined by plaque assays in Vero cells. For *in vitro* infections, cells were infected with HSV-1 or PRV-QXX at an MOI of 1. For *in vivo* studies, mice were intranasally inoculated with HSV-1 (1 × 10^6^ pfu/mouse).

### Immunoblotting, ubiquitination and Co-immunoprecipitation (Co-IP)

For immunoblotting, cells were lysed in RIPA buffer [50 mM Tris-HCl (pH 8.0), 150 mM NaCl, 1% Triton X-100, 1% sodium deoxycholate, 0.1% SDS, 2 mM MgCl_2_] supplemented with protease/phosphatase inhibitors. Proteins were resolved by SDS-PAGE, transferred to PVDF membranes, and blocked with 5% nonfat milk in TBST (1 hour, RT). Membranes were incubated with primary antibodies (overnight, 4°C), then HRP-conjugated secondary antibodies (1 hour, RT). Signals were detected using Laminate Crescendo Western HRP Substrate (Millipore, WBLUR0500) on a GE AI600 imager.

For ubiquitination assay, cells were lysed in IP buffer [50 mM Tris-HCl (pH 7.4), 150 mM NaCl, 1% NP-40, 1% sodium deoxycholate, 5 mM EDTA, 5 mM EGTA] and clarified (16,000 × *g*, 10 minutes, 4°C). Subsequently, 900 μL aliquots were incubated with 40 μL of a 1:1 slurry of Sepharose beads conjugated to anti-HDAC1 or anti-FLAG mouse monoclonal antibodies (4°C, 4 hours). Beads were washed four times with IP buffer, eluted in SDS sample buffer (10 minutes, boiling), and analyzed by immunoblotting.

For Co-IP assay, cells were lysed in Co-IP buffer [50 mM Tris-HCl (pH 7.4), 150 mM NaCl, 1% NP-40, 5 mM EDTA, 5 mM EGTA] and clarified (16,000 × *g*, 10 minutes, 4°C). Aliquots (900 μL) were incubated with 40 μL of a 1:1 slurry of Sepharose beads conjugated to IgG (GE Healthcare, AI600) or anti-FLAG mouse monoclonal antibody (4°C, 4 hours). Beads were washed three times with Co-IP buffer, eluted in SDS sample buffer (10 minutes, boiling), and analyzed by immunoblotting.

### Immunofluorescence assay

Cells were fixed in 4% paraformaldehyde at room temperature for 20 min on coverslips in 12-well plates. Following three washes with PBS, cells were permeabilized with 0.2% Triton X-100 for 20 min and subsequently blocked with 10% FBS. Specific primary antibodies, diluted in 10% FBS, were then applied to the cells and incubated for 1 h at room temperature. After three PBS washes, cells were exposed to the appropriate secondary antibodies, also diluted in 10% FBS, for 1 h at room temperature. Nuclei were stained with DAPI for 5 min at room temperature, mounted using Prolong Diamond (Invitrogen, P36970), and visualized using a Zeiss LSM 800 confocal microscope.

### Comet assay

Cells were seeded in 6-well plates and treated accordingly. Frosted microscope slides were coated with 0.5% normal melting point agarose. A mixture of 10 μL DMEM containing around 10,000 cells and 75 μL of 0.7% low melting point agarose was layered onto the pre-coated agarose. An additional 75 μL of 0.7% low melting point agarose formed a third layer. The slides were lysed in a buffer containing 10 mM Tris-HCl, pH 10.0, 2.5 M NaCl, 100 mM Na2EDTA, 1% Triton X-100, and 10% DMSO for 2 hours at 4°C. Post-lysis, the slides were treated with an electrophoresis solution (300 mM NaOH, 1 mM Na2EDTA, pH >13) for 40 minutes, electrophoresed at 20 V (~300 mA) for 25 minutes, and neutralized with 0.4 mM Tris-HCl (pH 7.5). Subsequently, cells were stained with propidium iodide (5 μg/mL) and examined using a Zeiss LSM 800 confocal microscope. DNA damage was assessed based on tail moment using CometScore software.

### qRT-PCR analysis

Total RNA was isolated using TRIzol reagent (TaKaRa, 9108) and reverse-transcribed into cDNA with the PrimeScript RT reagent kit (TaKaRa, RR047A). qRT-PCR was performed in triplicate using SYBR Premix Ex Taq (TaKaRa, RR820A) according to the manufacturer’s protocol. Expression levels were normalized to *β-actin* as an internal reference. Melting curve analysis confirmed amplification specificity by verifying single-product formation in all reactions. Relative gene expression was calculated using the 2^−ΔΔCt^ method. Primer sequences are provided in Table S1.

### Plaque assay

Vero cells (seeded in six-well plates) were grown to confluence, infected with serially diluted viruses (10^−1^–10^−7^; 1 hour, 37°C), and washed with PBS to remove residual inoculum. DMEM containing 1% methylcellulose (4 mL/well) was added, and cells were incubated for 4–5 days. Post-incubation, cells were fixed with 4% paraformaldehyde (15 minutes), stained with 1% crystal violet (30 minutes), and plaques were quantified.

### Statistical analysis

Statistical analyses were performed using GraphPad Prism 8 software. Comparisons between two groups were evaluated using a two-tailed Student’s *t*-test. Significance was defined as *P* < 0.05. Data are presented as mean ± SD of three independent experiments. Kaplan-Meier survival curves were generated and analyzed for mouse survival assessment.

## Results

### HSV-1 infection promotes HDAC1/2 degradation and histone hyperacetylation

Post-translational modifications of histones play a critical role in regulating chromatin remodeling and gene expression. HDACs, a highly conserved enzyme family [25], catalyze the removal of acetyl groups from histones, thereby promoting chromatin condensation and transcriptional repression. To investigate the effect of HSV-1 infection on HDAC expression and histone acetylation, we examined the protein levels of key HDAC isoforms following viral infection. Immunoblotting analysis revealed a marked decrease in HDAC1 and HDAC2 protein levels in cells infected with either HSV-1 or PRV, whereas the expression of HDAC4, HDAC6, and HDAC11 remained unchanged (Fig. 1A, B). In contrast, qRT-PCR results showed no alterations in HDAC1 and HDAC2 mRNA levels (Fig. 1C, D), suggesting that their depletion is regulated at the post-translational level.

**Figure 1.**
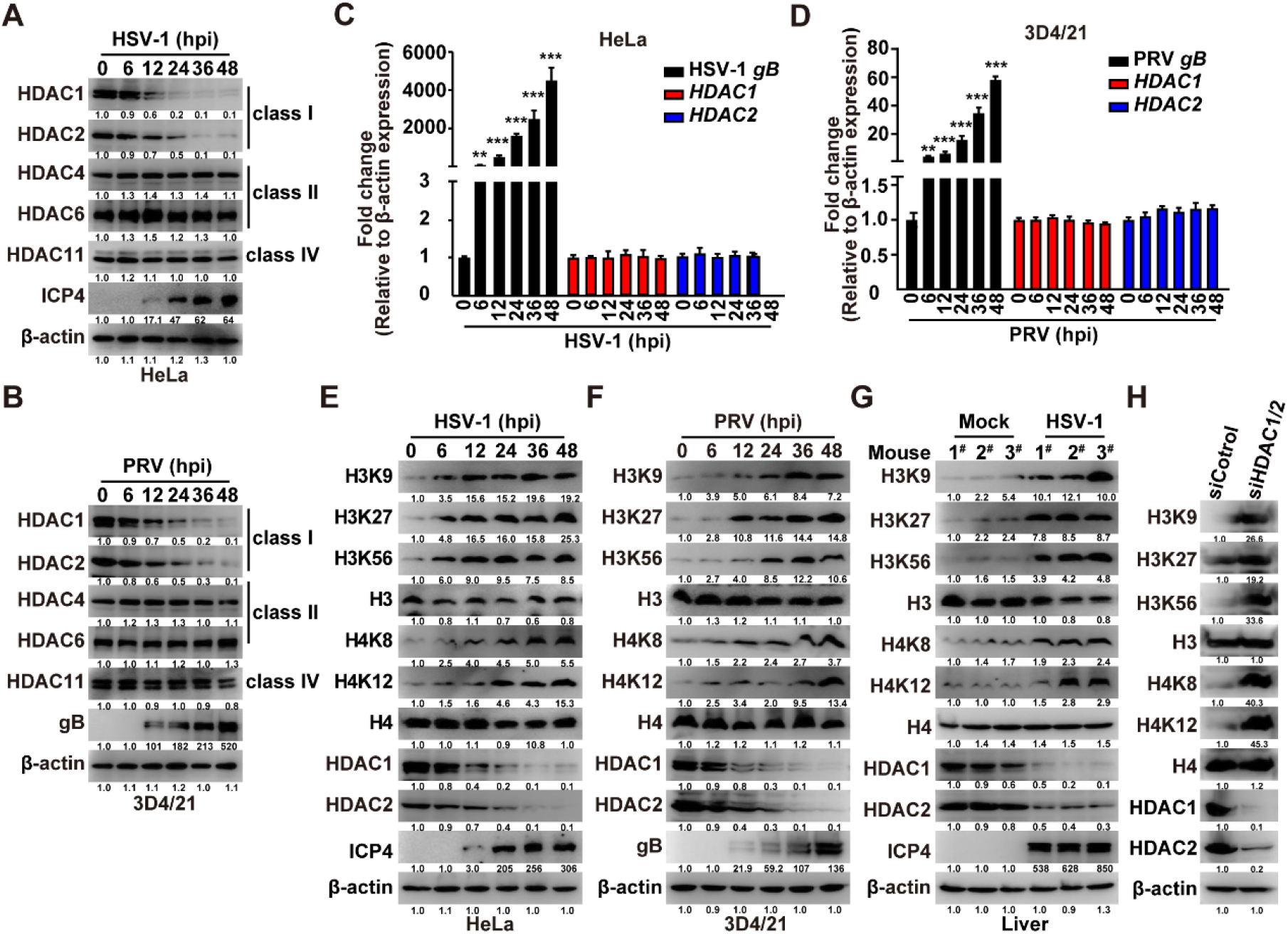
Viral infection induces degradation of class I HDACs and hyperacetylation of histones H3 and H4. (A) Immunoblotting analysis of the indicated HDACs in HeLa cells infected with HSV-1 (MOI = 1) for the indicated time points. (B) Immunoblotting analysis of the indicated HDACs in 3D4/21 cells infected with PRV-QXX (MOI = 1) for the indicated time points. (C and D) qRT-PCR analysis of *HDAC1* and *HDAC2* mRNA levels, normalized to β-actin expression, in HeLa cells infected with HSV-1 (C) or 3D4/21 cells infected with PRV-QXX (D) (MOI = 1). Data are mean ± SEM; n = 3. \*\**P* < 0.01, \*\*\**P* < 0.001. (E) Immunoblotting of the indicated proteins in HeLa cells infected with HSV-1 (MOI = 1) for the indicated time points. (F) Immunoblotting of the indicated proteins in 3D4/21 cells infected with PRV-QXX (MOI = 1) for the indicated time points. (G) Immunoblotting analysis of the indicated proteins in liver tissues from mice mock-infected or infected with HSV-1 (1 × 10^6^ pfu per mouse) at 5 days post-infection (n = 3). (H) Immunoblotting of the indicated proteins in HeLa cells transfected with siControl or siHDAC1/2. Note: The numerical values beneath each lane in the Western blot images represent the relative abundance of the corresponding protein bands, as quantified by densitometric analysis and normalized to the signal intensity of the first lane, which serves as the control group.

To assess the functional impact of HDAC1/2 reduction, we examined global histone acetylation patterns. Site-specific immunoblotting demonstrated elevated acetylation at multiple lysine residues, including H3K9, H3K27, H3K56, H4K8, and H4K12, in HSV-1-infected HeLa cells (Fig. 1E). A similar hyperacetylation profile was observed in PRV-infected 3D4/21 cells and in murine liver tissues infected with HSV-1 (Fig. 1F, G). Next, we assessed the effect of HDAC1/2 knockdown on global histone acetylation and observed significantly elevated acetylation at multiple evolutionarily conserved lysine residues–including H3K9, H3K27, H3K56, H4K8, and H4K12–consistent with genome-wide histone hyperacetylation (Fig. 1H). Collectively, these orthogonal lines of evidence—viral infection–induced degradation, cross-species conservation, and knockdown-mediated phenocopy—support the conclusion that HSV-1–mediated depletion of class I HDACs drives genome-wide histone hyperacetylation, establishing a chromatin environment permissive for viral transcription and replication.

### HSV-1 infection induces DDR through HDAC1/2 depletion and histone hyperacetylation

Given that excessive histone acetylation can destabilize chromatin structure, we sought to elucidate the mechanism by which this epigenetic modification activates the DDR pathway. Accumulating evidence underscores the pivotal role of histone modifications in maintaining genomic stability, with dynamic changes in chromatin architecture being intimately associated with DDR initiation. To determine whether HSV-1 infection induces DDR activation, we evaluated the phosphorylation of γ-H2AX, a well-established marker of DNA double-strand breaks. Immunofluorescence microscopy revealed a significant increase in γ-H2AX foci in cells infected with HSV-1 or PRV (Fig. 2A–C). Comet assays (single-cell gel electrophoresis) quantify DNA fragmentation, with an increased tail moment reflecting elevated levels of DNA damage–findings that further corroborate enhanced DNA fragmentation under these experimental conditions (Fig. 2D–G). Given the central role of ATM/ATR signaling in the DDR, we proceeded to examine the activation of key checkpoint kinases. Immunoblotting analysis confirmed phosphorylation of ATM, ATR, Chk1, Chk2, H2AX, and RAD51 in HSV-1-infected HeLa cells and 3D4/21 cells (Fig. 2H, I). Subsequently, we assessed the effect of HDAC1 knockdown on the DDR pathway and found that HDAC1-depleted HeLa cells exhibited reduced phosphorylation of ATM,ATR, and H2AX (Fig. 2J), indicating that HDAC1 depletion is sufficient to potentiate DDR signaling. Together, these data demonstrate that HSV-1 infection induces a coordinated, ATM/ATR-dependent DDR, and that HDAC1 loss represents a functionally relevant upstream event contributing to this response.

**Figure 2.**
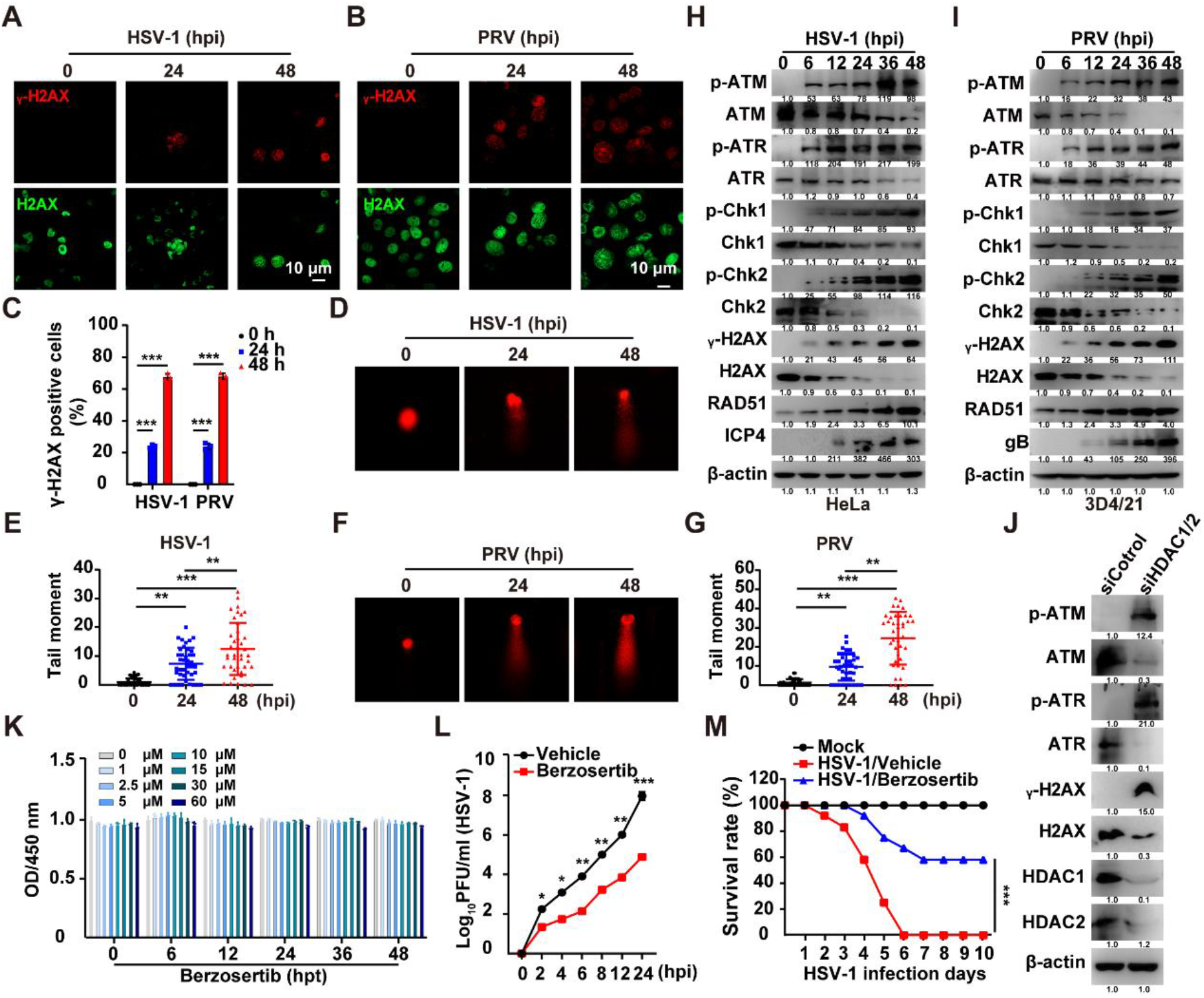
Viral infection triggers chromatin dysfunction and DDR activation via HDAC degradation. (A–C) Immunofluorescence staining of γ-H2AX (red) and H2AX (green) in HSV-1-infected HeLa cells (A) or PRV-infected 3D4/21 cells (B) (MOI = 1). Panel C presents the quantitative analysis of γ-H2AX foci intensity or nuclear fluorescence signal derived from panels A and B. Scale bars: 10 μm. Data are mean ± SEM; n = 100 cells/group. \*\*\**P* < 0.001. (D and E) Comet assay assessing DNA damage in HeLa cells infected with HSV-1 (MOI = 1). Data are mean ± SEM; n = 40 cells/group. \*\**P* < 0.01, \*\*\**P* < 0.001. (F and G) Comet assay assessing DNA damage in 3D4/21 cells infected with PRV (MOI = 1) at the indicated time points. Data are mean ± SEM; n = 40 cells/group. \*\**P* < 0.01, \*\*\**P* < 0.001. (H and I) Immunoblotting analysis of DDR markers (p-ATM, p-ATR, p-Chk1, p-Chk2, γ-H2AX) and viral ICP4/EP0 in HSV-1-infected HeLa cells (G) or PRV-QXX-infected 3D4/21 cells (H) (MOI = 1). (J) Immunoblotting analysis of DDR markers (p-ATM, p-ATR, γ-H2AX) and HDAC1/HDAC2 in HeLa cells transfected with siHDAC1/2. (K) HeLa cells were treated with vehicle and Berzosertib (0–60 μM) for 0–48 h. Cell proliferation was analyzed by CCK-8 assay. ns, no significance. (L) Viral titers in HSV-1-infected HeLa cells (MOI = 2) treated with berzosertib (50 nM) or vehicle. t = 0 h defined as time of medium replacement post-adsorption.Data are mean ± SEM; n = 3. \**P* < 0.05, \*\**P* < 0.01, \*\*\**P* < 0.001 (M) Survival curves of HSV-1-infected mice (1 × 10^6^ PFU/mouse) treated with berzosertib (20 mg/kg) or vehicle (n = 12/group), \*\*\**P* < 0.001. Note: The numerical values beneath each lane in the Western blot images represent the relative abundance of the corresponding protein bands, as quantified by densitometric analysis and normalized to the signal intensity of the first lane, which serves as the control group.

Next, we pharmacologically validated the functional relevance of ATR in HSV-1 replication using Berzosertib, a selective and clinically advanced ATR inhibitor. CCK-8 assays confirmed that the concentration of Berzosertib used (100 nM) did not significantly affect HeLa cell viability or proliferation over 48 hours (Fig. 2K), yet it markedly reduced viral titers in plaque assays (Fig. 2L). Collectively, these findings demonstrate that HSV-1 infection activates the DDR, likely through degradation of HDAC1 and HDAC2 and the resulting histone hyperacetylation. This virus-triggered DDR activation highlights the profound disruption of host chromatin homeostasis and genomic integrity by HSV-1.

### HSV-1 promotes the ubiquitin-proteasome degradation of HDAC1/2

To elucidate the mechanism responsible for HDAC1/2 degradation, we focused on their post-translational regulation. Since HDAC1/2 mRNA levels were unchanged after infection, we hypothesized that the reduction in HDAC1/2 protein is mediated through enhanced proteolytic degradation. To identify the relevant degradation pathway, HSV-1-infected HeLa cells were treated with either the proteasome inhibitor MG-132 or autophagy inhibitors (3-MA and chloroquine). Immunoblotting analysis indicated that only MG-132 rescued HDAC1/2 protein levels (Fig. 3A), confirming that degradation occurs primarily via the proteasomal pathway.

**Figure 3.**
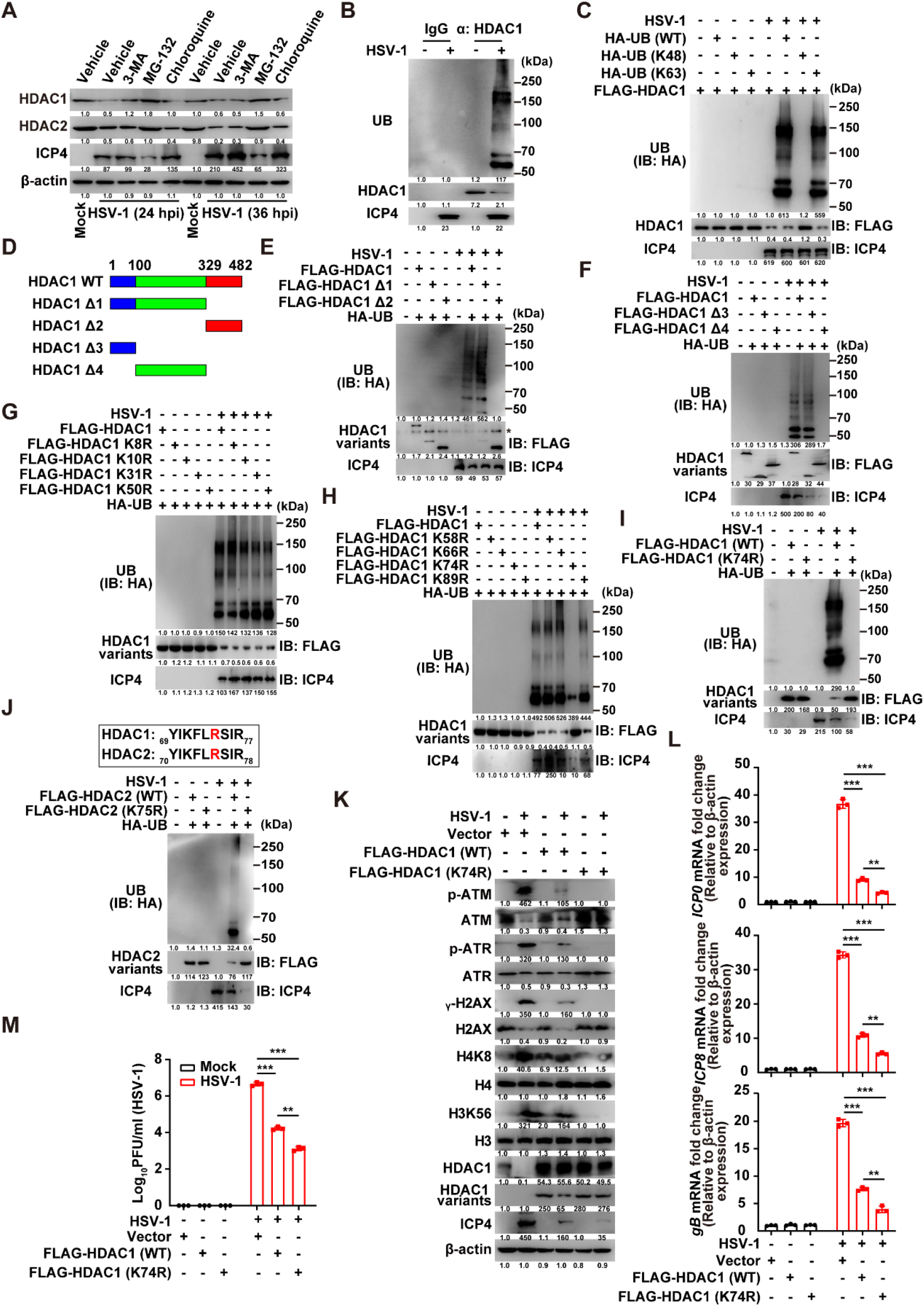
Viral infection induces K63-linked ubiquitination of HDAC1/2. (A) Immunoblotting analysis of the indicated proteins in HeLa cells either mock-infected or infected with HSV-1 (MOI = 1), treated with vehicle, 3-MA (10 μM), MG-132 (10 μM), or chloroquine (10 μM) for the indicated times. (B) Ubiquitination assay of HDAC1 in HeLa cells either mock-infected or infected with HSV-1 (MOI = 1) at 24 hpi. (C) Ubiquitination assay of FLAG-HDAC1 in HeLa cells transfected with HA-ubiquitin variants (WT, K48, K63), and either mock-infected or infected with HSV-1 (MOI = 1) at 24 hpi. (D) Schematic representation of HDAC1 deletion mutants. (E–I) Ubiquitination assays of FLAG-HDAC1 mutant variants in HeLa cells either mock-infected or infected with HSV-1 (MOI = 1) at 24 hpi. (J) Ubiquitination assays of FLAG-HDAC2 variants (WT and K75R) in HeLa cells either mock-infected or infected with HSV-1 (MOI = 1) at 24 hpi. (K) Immunoblotting analysis of the indicated proteins in HeLa cells transfected with empty vector, FLAG-HDAC1 (wild-type), or FLAG-HDAC1 (K74R), followed by mock-infected or infected with HSV-1 (MOI = 1) at 24 hpi. (L) qRT-PCR analysis of HSV-1 *ICP0, ICP8* and *gB* mRNA levels, normalized to β-actin expression, in HeLa cells transfected with empty vector, FLAG-HDAC1 (wild-type), or FLAG-HDAC1 (K74R), followed by mock-infected or infected with HSV-1 (MOI = 1) at 24 hpi. Data are mean ± SEM; n = 3. ns > 0.05, \**P* < 0.05, \*\**P* < 0.01,\*\*\**P* < 0.001. (M) Viral titer analysis in HeLa cells transfected with FLAG-HDAC1 or FLAG-HDAC1 (K74R), and either mock-infected or infected with HSV-1 (MOI = 1) at 24 hpi. Data are mean ± SEM; n = 3. \*\**P* < 0.01, \*\*\**P* < 0.001. Note: The numerical values beneath each lane in the Western blot images represent the relative abundance of the corresponding protein bands, as quantified by densitometric analysis and normalized to the signal intensity of the first lane, which serves as the control group.

To examine whether HSV-1 infection induced ubiquitination of HDAC1, endogenous immunoprecipitation assays were performed, which revealed increased ubiquitination of HDAC1 following infection (Fig. 3B). Subsequent ubiquitination assays using wild-type (WT), K48-only, and K63-only ubiquitin mutants demonstrated that HDAC1 undergoes K63-linked–but not K48-linked–polyubiquitination (Fig. 3C). Using a series of HDAC1 truncation mutants, we mapped the ubiquitination site to the N-terminal region (amino acids 1–100) (Fig. 3D–F). Site-directed mutagenesis further identified lysine 74 (K74) as the critical residue required for ubiquitination (Fig. 3G–I). Sequence alignment showed that HDAC2 K75 corresponds to HDAC1 K74 (Fig. 3J), and a K75R mutation in HDAC2 similarly abolished ubiquitination (Fig. 3J).

To further validate the role of HDAC1 K74, we first overexpressed HDAC1 WT and HDAC1 K74R and assessed phosphorylation of ATM, ATR, and H2AX; results showed that HDAC1 K74R overexpression suppressed HSV-1–induced DDR signaling (Fig. 3K). Furthermore, qRT-PCR analysis revealed that overexpression of both HDAC1 WT and HDAC1 K74R inhibited HSV-1 *ICP0, ICP8* and *gB* expression, but HDAC1 WT exerted a significantly stronger inhibitory effect (Fig. 3L). Consistently, HDAC1 K74R overexpression reduced viral titers in plaque assays (Fig. 3M). Collectively, these data demonstrate that HDAC1 K74 acetylation status modulates its capacity to restrain HSV-1 replication, likely by fine-tuning its regulatory influence on DDR signaling and viral gene expression

### MDM2 functions as the E3 ligase mediating HDAC1/2 ubiquitination and degradation

Having established the occurrence of K63-linked ubiquitination (Fig. 3), we next sought to identify the E3 ubiquitin ligase responsible. An RNAi screen targeting candidate E3 ligases revealed MDM2 as the principal mediator of HDAC1/2 degradation. Knockdown of MDM2–but not of PIRH2, KCTD11, CHFR, UHRF1, or TRIM46–effectively prevented HDAC1/2 depletion (Fig. 4A, B). Co-IP assay further confirmed an enhanced interaction between HDAC1 and MDM2 following HSV-1 infection (Fig. 4C).

**Figure 4.**
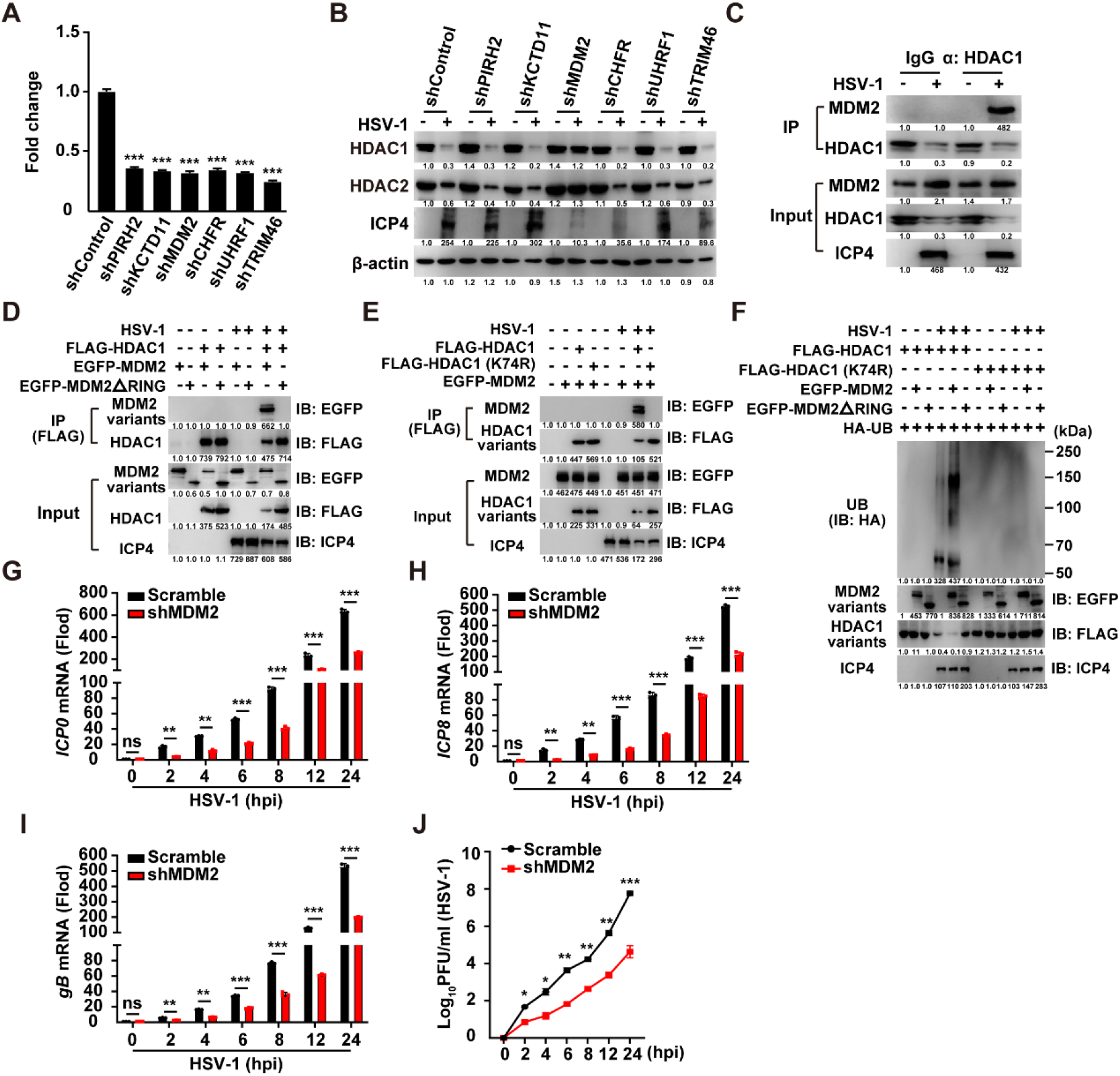
MDM2 mediates viral-induced HDAC1/2 degradation via K74/K75 ubiquitination. (A) qRT-PCR analysis validation of E3 ligase knockdown efficiency (shPIRH2, shKCTD11, shMDM2, shCHFR, shUHRF1, shTRIM46) in HeLa cells. Data are mean ± SEM; n = 3. \*\*\**P* < 0.001. (B) Immunoblotting analysis of HDAC1/2 stability in E3 ligase-knockdown HeLa cells infected with HSV-1 (MOI = 1). (C) Co-IP assay of endogenous HDAC1-MDM2 interaction in HSV-1-infected HeLa cells (MOI = 1; 24 hpi). (D) Co-IP assay of FLAG-HDAC1 with EGFP-MDM2 (WT or ΔRING) in HSV-1-infected HeLa cells (MOI = 1; 24 hpi). (E) Co-IP assay of FLAG-HDAC1 (WT or K74R) with EGFP-MDM2 in HSV-1-infected HeLa cells (MOI = 1; 24 hpi). (F) Ubiquitination assays of FLAG-HDAC1 (WT or K74R) co-expressed with EGFP-MDM2 (WT or ΔRING) in HSV-1-infected HeLa cells (MOI = 1; 24 hpi). (G–I) qRT-PCR analysis of HSV-1 *ICP0 (H), ICP8 (I)* and *gB (J)* mRNA levels, normalized to β-actin expression, in Scramble and shMDM2 HeLa cells, and either mock-infected or infected with HSV-1 (MOI = 1) at 24 hpi. Data are mean ± SEM; n = 3. ns > 0.05, \**P* < 0.05, \*\**P* < 0.01, \*\*\**P* < 0.001. (J) Viral titer analysis in Scramble and shMDM2 HeLa cells, either mock-infected or infected with HSV-1 (MOI = 2) at 24 hpi. t = 0 h defined as time of medium replacement post-adsorption. Data are mean ± SEM; n = 3. \*\**P* < 0.01, \*\*\**P* < 0.001. Note: The numerical values beneath each lane in the Western blot images represent the relative abundance of the corresponding protein bands, as quantified by densitometric analysis and normalized to the signal intensity of the first lane, which serves as the control group.

To characterize this interaction, we focused on the RING finger domain of MDM2, which is essential for its E3 ligase activity. Co-IP analysis demonstrated that HSV-1 infection promoted binding between FLAG-HDAC1 and EGFP-MDM2, but not with a RING-deleted mutant (EGFP-MDM2ΔRING) (Fig. 4D), indicating that the interaction was RING domain-dependent. To identify the region within HDAC1 required for binding, HeLa cells were co-transfected with EGFP-MDM2 and either FLAG-HDAC1 or the ubiquitination-resistant mutant FLAG-HDAC1 K74R. Infection with HSV-1 failed to induce binding between EGFP-MDM2 and the K74R mutant (Fig. 4E), underscoring the essential role of lysine 74 in mediating MDM2-dependent ubiquitination.

Ubiquitination assay further revealed that overexpression of EGFP-MDM2ΔRING did not enhance HSV-1-induced ubiquitination or degradation of HDAC1 (Fig. 4F). Similarly, the FLAG-HDAC1 K74R mutant remained resistant to ubiquitination under all experimental conditions (Fig. 4F). To further validate the role of MDM2, we performed MDM2 knockdown in infected cells; qRT-PCR analysis showed that MDM2 knockdown significantly suppressed HSV-1 *ICP0, ICP8*, and *gB* mRNA transcription (Fig. 4G–I), and reduced viral titers in plaque assays (Fig. 4J). Collectively, these results demonstrate that HSV-1 induces HDAC1/2 degradation through MDM2-mediated K63-linked ubiquitination.

### HSV-1 triggers cytoplasmic translocation of HDAC1 to facilitate its degradation

To elucidate the mechanisms by which HSV-1 induces HDAC degradation, we systematically analyzed the subcellular localization of HDAC1. In mock-infected HeLa cells, HDAC1 was predominantly nuclear; however, HSV-1 infection triggered a pronounced translocation of HDAC1 to the cytoplasm (Fig. 5B). This redistribution was completely abolished by treatment with leptomycin B (LMB), a specific inhibitor of CRM1-mediated nuclear export (Fig. 5B). At the same time, CCK-8 assay showed that LMB had no significant effect on cell proliferation (Fig. 5A).

**Figure 5.**
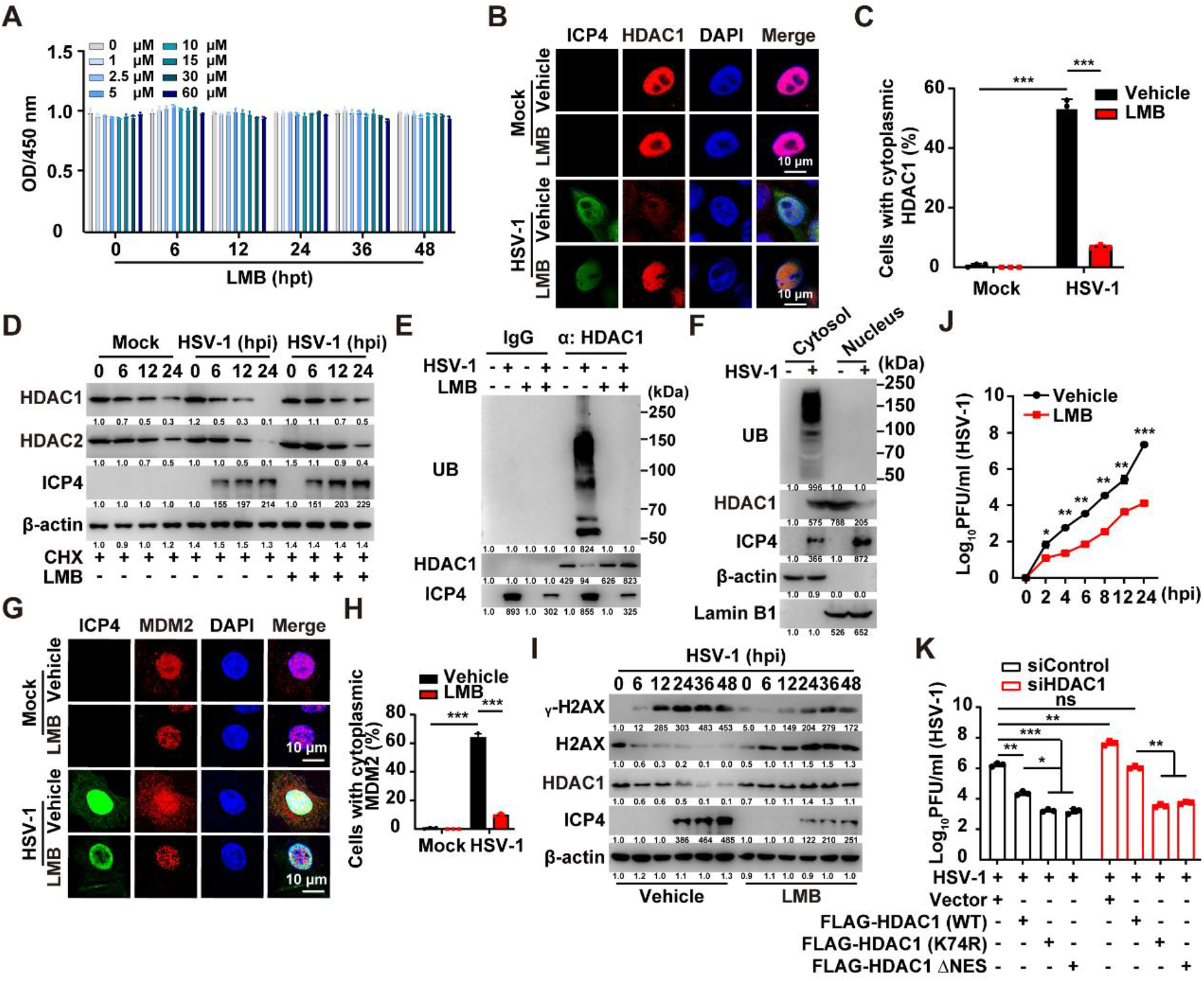
Cytoplasmic translocation enables MDM2-mediated HDAC1 degradation. (A) HeLa cells were treated with vehicle and LMB (0–60 μM) for 0–48 h. Cell proliferation was analyzed by CCK-8 assay. ns, no significance. (B and C) Immunofluorescence staining of HDAC1 (red) and viral ICP4 (green) in HSV-1-infected HeLa cells (MOI = 1) ± leptomycin B (LMB; 10 nM; 24 h). Scale bar: 10 μm. (C) shows the quantitative analysis of (B), Data are mean ± SEM; n = 100. \*\*\**P* < 0.001. (D) CHX chase assay measuring HDAC1/2 degradation kinetics in HSV-1-infected HeLa cells (MOI = 1) ± LMB (10 μM). (E) Ubiquitination assay of HDAC1 in HSV-1-infected HeLa cells (MOI = 1) ± LMB (10 μM; 24 h). (F) Immunoblotting analysis of ubiquitination in HDAC1 from cytosolic and nuclear fractions of HeLa cells infected with HSV-1. (G and H) Immunofluorescence analysis of MDM2 (red) and ICP4 (green) in HSV-1-infected HeLa cells (MOI = 1) ± LMB (10 μM; 24 h). Scale bar: 10 μm. (H) shows the quantitative analysis of (G), Data are mean ± SEM; n = 100. \*\*\**P* < 0.001. (I) Immunoblotting analysis of DDR markers (γ-H2AX) and HDAC1/2 in HSV-1-infected HeLa cells (MOI = 1) ± LMB (10 μM). (J) Viral titers in HSV-1-infected HeLa cells (MOI = 2) ± LMB (10 μM). t = 0 h defined as time of medium replacement post-adsorption.Data are mean ± SEM; n = 3. \**P* < 0.05, \*\**P* < 0.01, \*\*\**P* < 0.001 (K) Viral titer analysis in HeLa cells transfected with siHDAC1 transfected with empty vector, FLAG-HDAC1, FLAG-HDAC1 (K74R) or FLAG-HDAC1 ΔNES, infected with HSV-1 (MOI = 1) at 24 hpi. Data are mean ± SEM; n = 3. \*\**P* < 0.01, \*\*\**P* < 0.001. Note: The numerical values beneath each lane in the Western blot images represent the relative abundance of the corresponding protein bands, as quantified by densitometric analysis and normalized to the signal intensity of the first lane, which serves as the control group.

To determine whether translocation is functionally linked to degradation, we conducted cycloheximide (CHX) chase assays. Degradation of both HDAC1 and HDAC2 began approximately 12 hours after CHX treatment and was markedly accelerated by HSV-1 infection (Fig. 5D). Importantly, LMB treatment restored degradation kinetics to levels comparable to those in mock-infected cells (Fig. 5B). Moreover, under LMB treatment, HSV-1 was unable to induce ubiquitination of endogenous HDAC1 (Fig. 5E). To further validate the subcellular site of HDAC1 ubiquitination, we performed nucleocytoplasmic fractionation followed by immunoblotting; results showed that HSV-1–induced ubiquitination of endogenous HDAC1 occurs specifically in the cytoplasm (Fig. 5F).

We further demonstrated that nuclear export and subsequent proteasomal degradation of HDAC1 are strictly dependent on the E3 ubiquitin ligase MDM2. HSV-1 infection induces CRM1-dependent ubiquitination of endogenous HDAC1, thereby establishing CRM1-mediated nuclear export as an essential prerequisite for MDM2-catalyzed HDAC1 degradation (Fig. 5G, H). Consistent with this, immunoblotting analysis revealed that LMB significantly suppressed HSV-1–induced phosphorylation of p53 and γH2AX (Fig. 5I). Notably, LMB treatment-by inhibiting CRM1-dependent nuclear export and thereby preventing HDAC1 degradation–markedly impaired HSV-1 replication (Fig. 5F). To rigorously test whether HDAC1 nuclear export is the critical, rate-limiting step enabling its ubiquitination and degradation, we employed NESmapper to identify a canonical nuclear export signal (NES) in HDAC1 (residues 207–216). We then generated an HDAC1 ΔNES deletion mutant and performed functional rescue experiments in HDAC1-knockdown cells by reconstituting expression of HDAC1 WT, the ubiquitination-deficient mutant HDAC1 K74R, or FLAG-tagged HDAC1 ΔNES. Only HDAC1 WT fully restored the pro-viral phenotype associated with HDAC1 depletion; in contrast, neither HDAC1 K74R nor HDAC1 ΔNES rescued viral replication, and both exerted potent antiviral activity–significantly suppressing HSV-1 proliferation relative to controls (Fig. 5K). Collectively, these findings establish that HSV-1 co-opts the CRM1 nuclear export machinery to translocate HDAC1 to the cytoplasm, where it becomes accessible to MDM2 for ubiquitination and subsequent proteasomal degradation, thereby modulating host antiviral defense.

## Discussion

HSV-1 manipulates host cellular pathways to facilitate its replication, latency, and reactivation [26]. In this study, we demonstrate that alphaherpesvirus targets class I histone deacetylases HDAC1 and HDAC2 for MDM2-mediated K63-linked polyubiquitination and subsequent proteasomal degradation. This degradation induces histone hyperacetylation, chromatin relaxation, and enhanced viral replication–revealing a mechanism through which alphaherpesvirus reprograms the host epigenetic landscape to optimize viral replication (Fig. 6).

**Figure 6.**
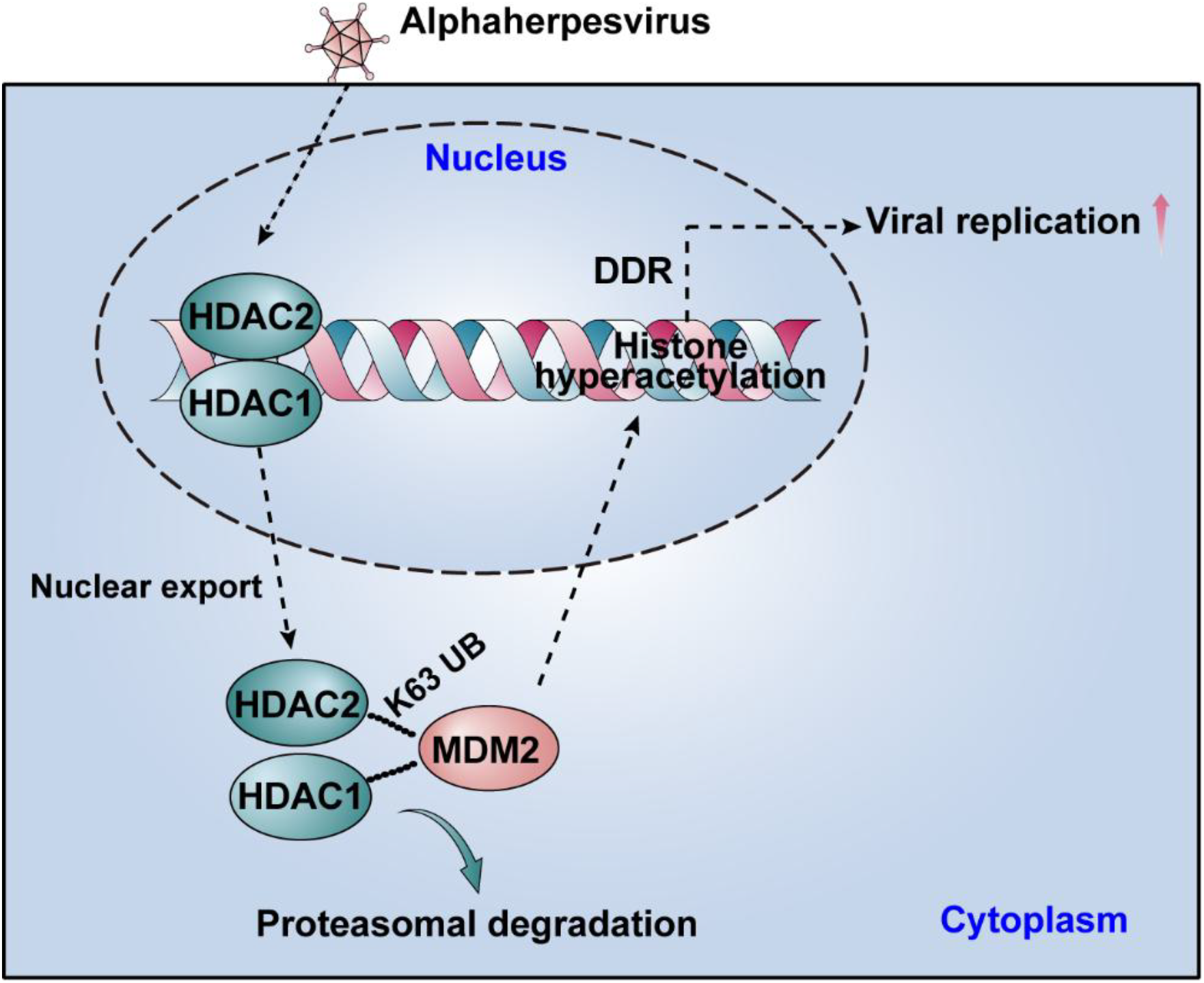
A schematic model showing the targeted cytosolic degradation of class I histone deacetylases is essential for efficient alphaherpesvirus replication.

In contrast, HSV-1 actively targets HDAC1 and HDAC2 for proteasomal degradation to facilitate lytic replication–a strategy that stands in marked contrast to human cytomegalovirus (HCMV), which relies on HDAC activity to maintain latency [27]. This divergence underscores a fundamental difference in epigenetic regulation among herpesviruses. Our data demonstrate that HSV-1–mediated HDAC1/2 degradation initiates as early as 4–6 hours post-infection, coinciding with the onset of immediate-early gene expression and preceding peak viral DNA synthesis. These findings support a context-dependent, dual functional role for HDAC1/2: their enzymatic activity helps enforce transcriptional silencing during latency, whereas their timely removal is essential for chromatin remodeling and efficient lytic gene expression.

We identified MDM2 [28] as the principal E3 ubiquitin ligase responsible for ubiquitinating HDAC1 at lysine 74 and HDAC2 at lysine 75, leading to their proteasomal degradation. This mechanism is distinct from other E3 ligases, including PIRH2 [29] and TRIM46 [30], and critically depends on the RING domain of MDM2 [31]. While K63-linked ubiquitination is typically associated with non-proteolytic functions, our work establishes its novel role in mediating HDAC1/2 degradation–uncovering a previously unrecognized viral pathogenesis pathway. Targeting this axis using MDM2 inhibitors or inhibiting nuclear export of HDAC1/2 significantly restricts viral replication, suggesting that host-directed antiviral strategies may help overcome resistance to direct-acting agents [31].

Depletion of HDAC1/2 increases acetylation at histone residues associated with open chromatin and active transcription. Similar chromatin remodeling has been observed in other viral infections [32], underscoring its conserved role in viral gene regulation. Our results are consistent across multiple cell types, including HeLa, 3D4/21, and murine liver, emphasizing the central importance of HDAC1/2 suppression in HSV-1 biology. Notably, HDAC1/2 degradation activates DDR, as indicated by increased γ-H2AX foci and phosphorylation of ATM, ATR, Chk1, and Chk2. This aligns with previous reports that HSV-1 exploits DDR signaling to enhance replication [33-35]. Importantly, HDAC1/2 undergo CRM1-dependent nuclear export prior to degradation–a process inhibited by leptomycin B–highlighting the essential role of subcellular localization in viral manipulation of host proteins.

HSV-1–mediated HDAC1/2 degradation likely supports viral replication through multiple mechanisms, including evasion of nuclear antiviral sensors and stimulation of viral gene expression. Targeting the MDM2–HDAC1/2 axis represents a promising antiviral strategy. MDM2 inhibitors reduce HSV-1 titers [36], suggesting that modulating ubiquitination pathways could complement existing therapies. Beyond herpesviruses, the MDM2–HDAC axis may be exploited by other DNA viruses that manipulate chromatin and DDR [32], indicating a possible common host vulnerability. However, broad HDAC inhibition may unintentionally enhance viral replication, underscoring the need for targeted therapeutic approaches [37].

This study primarily employed *in vitro* models (HeLa cells), which lack the neuronal environment essential for studying HSV-1 latency. Future work should validate these findings in primary neurons and trigeminal ganglia [38]. Additional studies are also needed to determine whether HDAC1/2 degradation occurs during reactivation or is limited to lytic infection [7], and to explore the interplay between HDAC degradation, viral tegument proteins, and host stress responses. *In vivo* evaluation of inhibitors targeting the MDM2–HDAC1/2 pathway will be crucial for assessing their therapeutic potential.

## Author contributions

Sheng-Li Ming and Meng-Hua Du: Data curation, Formal analysis, Investigation, Methodology, Validation, Visualization, Writing – original draft; Jia-Ming Yang: Data curation, Formal analysis, Investigation, Validation; Ya-Di Guo: Data curation, Formal analysis, Investigation, Validation, Visualization; Jia-Jia Pan: Data curation, Formal analysis, Investigation, Methodology, Visualization; Wei-Fei Lu: Data curation, Formal analysis, Investigation; Jiang Wang: Methodology, Visualization, Project administration, Supervision, Writing – review & editing; Lei Zeng and Bei-Bei Chu: Conceptualization, Data curation, Funding acquisition, Investigation, Methodology, Project administration, Resources, Supervision, Validation, Writing – original draft, Writing – review & editing.

## Ethics statement

All procedures involving animals in this study received ethical approval from the Institutional Animal Care and Use Committee (IACUC) of Henan Agricultural University (Permit Number: HNND2020031008). This research was performed in accordance with the regulations specified in the Guide for the Care and Use of Laboratory Animals established by the Ministry of Science and Technology of China

## Data Availability Statement

The datas that support the findings of this study are openly available in “Mendeley Data, V3” at https://data.mendeley.com/datasets/yg5fgtvxzk/3, doi: 10.17632/yg5fgtvxzk.3, reference number[39].

## Declaration of generative AI and AI-assisted technologies in the writing process

This study affirms that neither generative artificial intelligence nor any other AI-assisted technologies were employed in the preparation of this manuscript.

## Figure legends

**Table S1.**
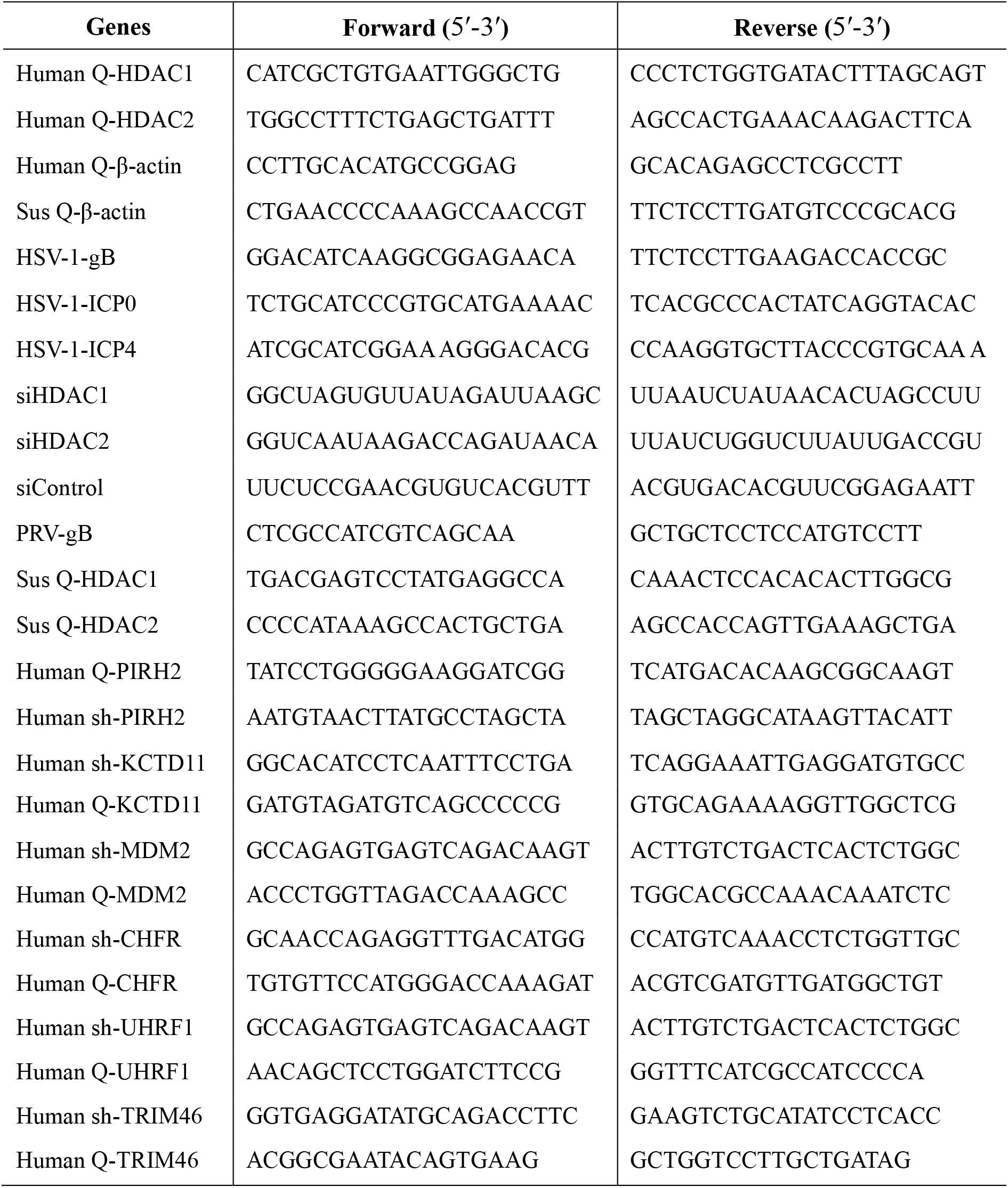
List of shRNAs and primers used in this study.

